# Learning attention-controllable border-ownership for objectness inference and binding

**DOI:** 10.1101/2020.12.31.424926

**Authors:** Antoine Dedieu, Rajeev V. Rikhye, Miguel Lázaro-Gredilla, Dileep George

## Abstract

Human visual systems can parse a scene composed of novel objects and infer their surfaces and occlusion relationships without relying on object-specific shapes or textures. Perceptual grouping can bind together spatially disjoint entities to unite them as one object even when the object is entirely novel, and bind other perceptual properties like color and texture to that object using object-based attention. Border-ownership assignment, the assignment of perceived occlusion boundaries to specific perceived surfaces, is an intermediate representation in the mammalian visual system that facilitates this perceptual grouping. Since objects in a scene can be entirely novel, inferring border ownership requires integrating global figural information, while dynamically postulating what the figure is, a chicken-and egg process that is complicated further by missing or conflicting local evidence regarding the presence of boundaries. Based on neuroscience observations, we introduce a model – the cloned Markov random field (CMRF)– that can learn attention-controllable representations for border-ownership. Higher-order contour representations that distinguish border-ownerships emerge as part of learning in this model. When tested with a cluttered scene of novel 2D objects with noisy contour-only evidence, the CMRF model is able to perceptually group them, despite clutter and missing edges. Moreover, the CMRF is able to use occlusion cues to bind disconnected surface elements of novel objects into coherent objects, and able to use top-down attention to assign border ownership to overlapping objects. Our work is a step towards dynamic binding of surface elements into objects, a capability that is crucial for intelligent agents to interact with the world and to form entity-based abstractions.

## Introduction

Human visual systems can parse a scene composed of novel objects and infer their surfaces and occlusion relationships without relying on object identities, or category-identifying textures [1](Fig 1A). This process involves perceptual grouping [2] whereby contour and surface elements belonging to the same object are linked and grouped, permitting them to be separated from other scene components. Perceptual grouping can bind together spatially disjoint entities to unite them as one object even when the object is entirely novel, and bind [3] other perceptual properties like color and texture to that object using top-down object-based attention. Flexible deployment of this ability is crucial for interacting with the world and for formation of perceptually grounded higher-level abstractions [4, 5, 6].

**Figure 1:**
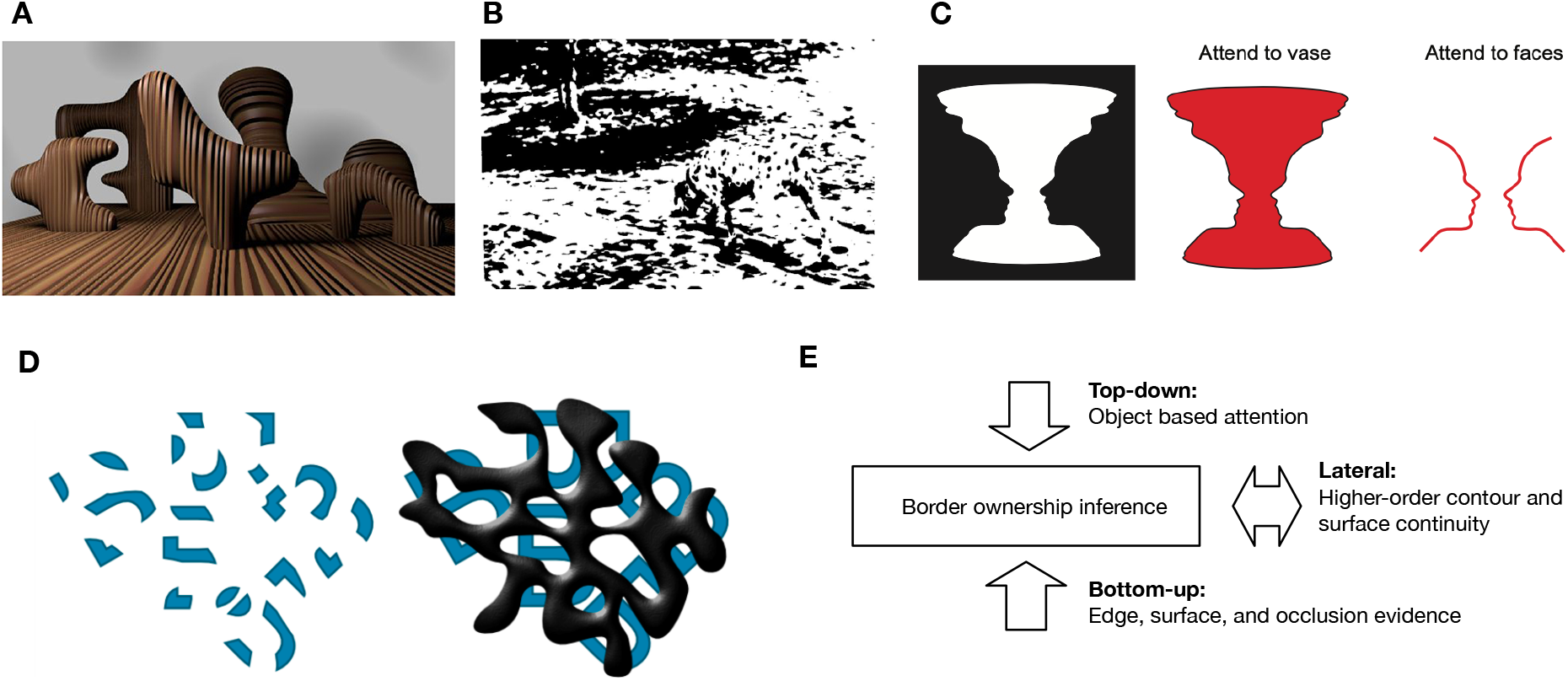
Examples of objectness and border-ownership inference and binding. **A**: People can parse visual scenes consisting of entirely novel objects, infer their borders and surfaces, and reason about occlusions, tasks that are challenging for current AI systems. Figure credit [1] **B**: A challenging example of visual inference where the inferring the boundaries of dalmation dog is tightly coupled with the perception of the dog itself. **C**: Rubin’s vase illusion is a commonly used example for multi-modality of border ownership, as depending on the focus of attention, one can perceive either a vase or two faces, and the ownership assignment of the same border changes accordingly. **D**: Bregman’s Bs [16] are a famous illustration of the relationship between binding, occlusions, and border-ownership inference. In the first panel, the different surface segments are not easily perceived and bound together as instances of the letter B, whereas in the second panel the very same surface segments are bound together as unitary objects. **E**: Border-ownership inference requires systematically integrating bottom-up, top-down and lateral evidence

In mammalian vision, perceptual grouping of novel objects is known to be facilitated via an intermediate representation of border-ownership [7, 8, 9, 10]. Border-ownership refers to the assignment of occlusion boundaries in images to object percepts, a chicken-and-egg problem because the object percepts themselves depend on the assignment of borders and surfaces [11]. Since objects in the scene can be entirely novel, inferring border ownership requires integrating global figural information, while dynamically postulating what the figure is, a process that is complicated further by missing or conflicting local evidence regarding the presence of boundaries Fig 1B. While contour-integration is known to be part of this process, the underlying model will need to account for higher-order relationships [12, 13]between contour elements in a scene, not just pair-wise. Border ownership is also multistable [14, 15]since the same border can be assigned different ownership based own top-down attention as shown by the famous face-vase phenomenon Fig 1C. Moreover, the presence or absence of occlusion cues significantly affects the grouping process, and the assignment of border ownership. In the famous Bregman’s Bs effect [16], the segments of B’s are not perceived as unitary objects without an occlusion cue (Fig 1D first panel), while the presence of an occlusion cue aids the perceptual grouping of the same disconnected surface segments (Fig 1D second panel). These examples, and evidence from neuroscience and cognitive science [7, 10, 17, 11, 18, 19, 9, 20] suggest that border ownership computation requires learning representations that allow for dynamic integration of bottom-up, lateral, and top-down information in a systematic manner (Fig 1E).

Numerous computational models, emphasizing different aspect of border-ownership inference, have been developed by both computer vision and computational neuroscientists. Computer vision approaches generally attempt to solve the instance segmentation problem by providing higher levels of features, such as object class. Inferring figure-ground organization in two-dimensional (2D) static images have been either addressed by limiting the input to line drawings [21], or by extracting local features for natural images and applying machine learning techniques [22, 23]. Usually, approaches in the latter category formulate the problem in a conditional random field (CRF) framework to incorporate context into their model. A number of these models, such as [24], employ shape-related features like convexity-concavity of the contour segment or local geometric classes, such as horizontal features (sky, ground, etc.), or vertical features (buildings, etc.). Recently, a CNN-based model, called DOC, for figure-ground organization in a single 2D image has been introduced [23]. This network consists of two streams, one detecting edges and the other specifying border ownership, solving a joint labeling problem. The feed-forward network, while having the advantage of incorporating diverse cues to predict occlusion boundaries, is restricted to unimodal predictions, whereas border owneership predictions can be influenced by context and top-down attention. Computational neuroscience models [25, 26], on the other hand, use explicitly constructed features and lateral interactions to model different aspects of border-ownership, but often does not learn the representations and are not tested on a wide range of challenging inputs. Several recent neural network models have recognized the importance of long-range lateral connections [13, 27]in contour-detection, but are yet to tackle the interconnected challenges of multi-stable border-ownership, closure, occlusion, and objectness.

In this paper, we present an undirected probabilistic graphical model (PGM) that learns border-ownership and surface representations such that the hidden higher-order border-ownership assignments can be inferred from noisy and occluded test inputs, and controlled using top-down attention. Neuroscience observations and earlier computational models guided the scaffolding of the model that allows it to learn such representations. Our model takes the form of a cloned Markov random field (CMRF) and is a natural, two-dimensional extension of the recently developed cloned hidden Markov model [28], and clone-structured cognitive graphs [29]. A key insight in this representation is *clonal* states [30] that represent the same bottom-up input, but learn to wire to different contexts such that higher-order contour contexts can be represented efficiently. In the CMRF, the hidden variables form an eight-connected grid, with each of them being connected to an observation, which is an image pixel. With this wiring, the different clones have the potential to learn to represent the different higher-order contours, border-ownership assignments, or surface patches a given pixel might belong to. To train the CMRF we utilized a novel approach, query training [31], to turn the CMRF into a trainable inference neural network. We show that higher-order contour representations, and border-ownership-signaling neurons emerge as part of learning in this CMRF, and that their inference dynamics are similar to those observed in neurophysiology. When tested with a cluttered scene of novel 2D objects rendered only using noisy contour-evidence, the CMRF model is able to perceptually group them, despite clutter and missing edges. Moreover, the CMRF is able to use occlusion cues to bind disconnected surface elements of novel objects into coherent objects, and able to use top-down attention to assign border ownership to overlapping objects.

We structure our paper as follows. First, we discuss the model in detail. Next, we demonstrate the performance of the model on several proposed datasets as well as its ability to mimic known psychophysical phenomena. Finally, we discuss how our model differs from existing models of border ownership.

## Results

### Cloned Markov random field (CMRF) as a surface model

A Markov random field (MRF) is a set of random variables that follow a probability distribution associated with an undirected graph *G*. If we call these variables 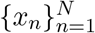, the probability distribution factorizes over the cliques of *G* as 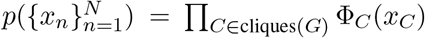, where Φ_*C*_ (·) is the factor potential associated to clique *C* and *x*_*C*_ the subset of variables in *C*. In a grid MRF the variables are arranged forming a 2D grid and the graph *G* connects each variable to the adjacent ones. When each variable is connected specifically to the 8 variables immediately surrounding it in the grid, the resulting probability distribution is called an 8-connected grid MRF. In general, these variables could be discrete or continuous, but we only consider the discrete case in this work.

Grid MRFs are often used in image processing applications (Li, 2009), but the MRF variables (which we will call “pixel labels”) are always observed when learning the factor parameters. We propose herein a model, referred to as cloned Markov random field (CMRF), whose structure allows learning higher-order contours, contour-surface interactions, and border-ownership. During inference, the model can perceptually group disconnected fore-ground elements, separate them from cluttered backgrounds and assign border ownerships. An example of a CMRF is presented in **Fig. 2B**, and introduced below. In contrast to grid MRFs, the pixel labels in this model are categorical *hidden* variables, i.e., we do not have access to the labels for training. Our hidden variables possess two types of uncertainty: (1) categorical (ternary) *pixel labels* are combined into binary *pixel intensities* through a noisy channel, which prevents us from knowing the exact labels; and (2) each pixel label has multiple *clones* associated with it, with each clone deterministically emitting the same pixel label. Consequently, the problem of recovering the active clone associated with an active label is non-trivial.

**Figure 2:**
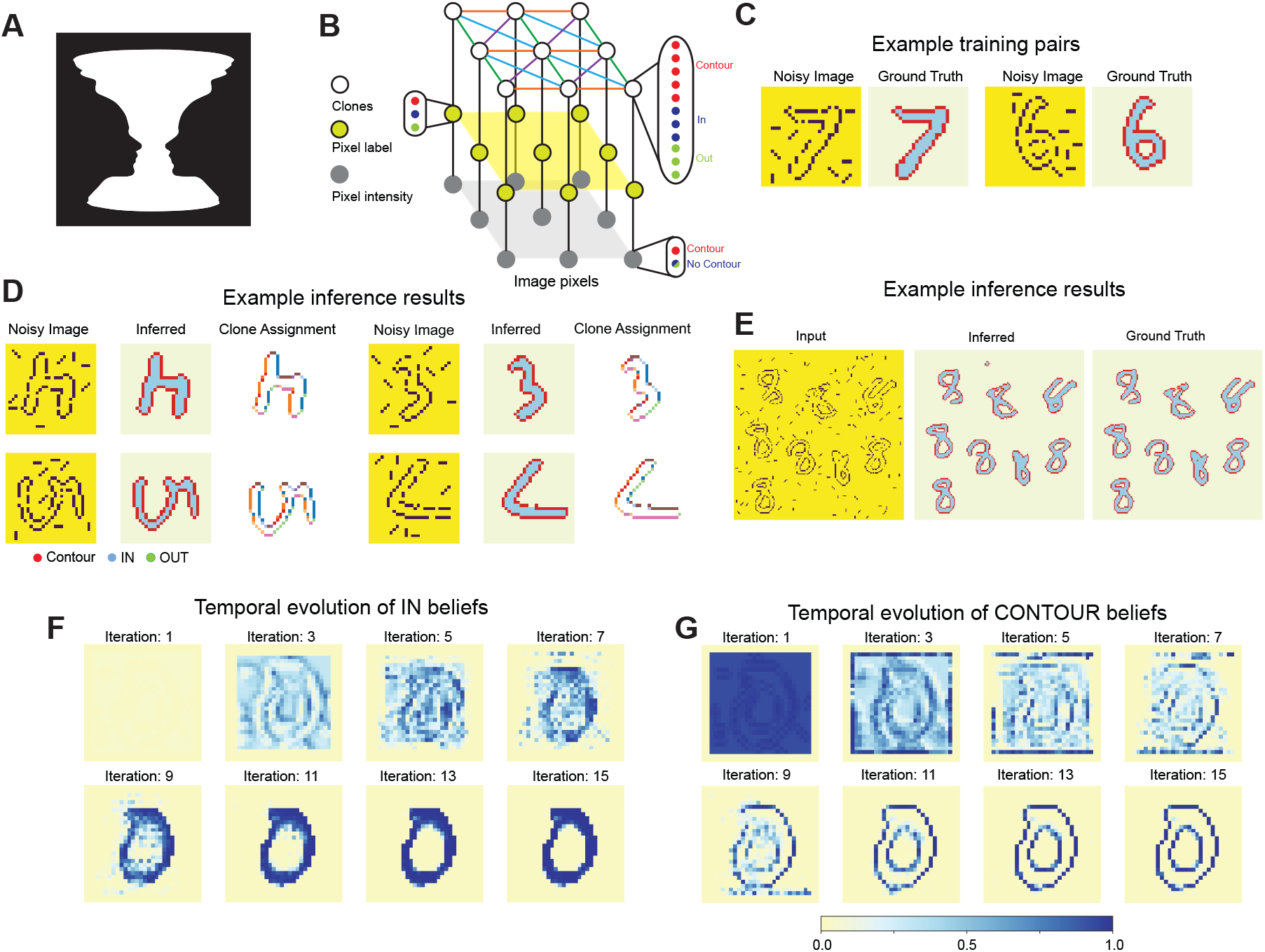
CMRFs can be learned using query training (QT) and perform foreground-background segmentation of new unseen digits. **A**: Rubin’s vase illusion is a commonly used motivating example for border ownership, as depending on the focus of attention, one can perceive either a vase or two faces. **B**: Schematic of cloned Markov Random Field used as a surface model. In this model, categorical hidden clones (11 categories) deterministically emit ternary (`IN, OUT` and `CONTOUR`) pixel labels. In turn, each pixel label emits an observed binary pixel intensity (0 or 1) through a noisy channel. **C**: Example input-output pairs of the first part of the training set used to train the CMRF with Query Training. Each noisy binary input is paired with a ground truth segmentation of `IN`, `OUT` and `CONTOUR` pixel labels. **D**: Test images examples: 20% of the `CONTOUR` pixels are missing and spurious edges are added. Despite these perturbations, the CMRF is still able to accurately segment the foreground from the background. Clones capture higher order properties of the contour. **E**: The CMRF can segment an image of 8s digits which are unseen during training and upsampled. **F-G**: Temporal evolution of `IN` and `CONTOUR` beliefs for a test example digit. See also Supplementary **Fig. 4** and Supplementary Videos 1 and 2.

The central idea behind cloned Markov random fields is dynamic Markov coding [32], which is a method for representing higher-order sequences by splitting, or cloning, observed states. For example, a first-order Markov chain representing the sequence of events *A* − *C* − *E* and *B* − *C* − *D* will assign a 0.25 probability to the sequence *A* − *C* − *D*. In contrast, dynamic Markov coding makes a higher-order model by splitting the state representing event *C* into two copies, one for each incoming connection: the first copy *C*_1_ specifies the sequence *A* − *C*_1_ − *E*, while the second copy *C*_2_ represents the sequence *B* − *C*_2_ − *D*. This state cloning mechanism permits a sparse representation of higher-order dependencies and has been discovered in various domains [33, 34, 35, 32]. With cloning, the same bottom-up sensory input (*C* in our example) is represented by a multitude of states (*C*_1_ and *C*_2_ in our example) that are copies of each other in their selectivity for the sensory input, but specialized for specific temporal contexts, enabling the efficient storage of a large number of higher-order and stochastic sequences without destructive interference. The CMRF that we present in this paper is a natural extension of the sequential analog presented in [28, 29].

The central idea in CMRFs of using different copies of the same observation for different higher-order contexts is similar to the proposal in [36, 37]for the computational role of clonal neurons in a cortical column [38]. The CMRF can be readily instantiated as a neuronal circuit similar to the ones proposed in [36, 29]. Each clone corresponds to a neuron, and the *lateral connections* between these neurons make up the transition matrix. All clones receive the same bottom-up or feedforward input from the observation, and they learn different representations by wiring to different lateral contexts. The output of these clones is then a weighted sum of its lateral inputs, multiplied by the bottom-up input [39, 29].

A simplified version of our CMRF, based on these principles is presented in **Fig. 2B**. This model has three types of variables: (1) clones (white dots), which are hidden categorical variables (11-dimension in the figure), (2) pixel labels (yellow dots), which are categorical ternary hidden variables, and (3) pixel intensities (gray dots), which are observed binary variables. Pixel labels indicate a pixel as belonging to the `OUT` (green dots), `IN` (blue dots), or `CONTOUR` (red dots) of the foreground surface. Clones deterministically emit pixel labels, with multiple clones emitting the same pixel label. In particular, in the model presented in **Fig. 2B**, the `OUT, IN` and `CONTOUR` labels are respectively emitted by three, three and five different clones. By having five apparently identical clones, all of which generate a `CONTOUR` label, the CMRF model can specialize and learn higher-order properties of the contour, such as its orientation and the side of the foreground on which it is located. For example, a pixel that is part of a horizontal contour will have a different active clone than another pixel that is part of a vertical contour. Another advantage of this model structure is that the context captured by the active clone of a pixel gives information about its neighboring pixels. For instance, a `CONTOUR` pixel label whose active clone captures a “bottom-horizontal” orientation is likely to turn on the same “bottom-horizontal” clone on its left and right and a clone corresponding to the `IN` pixel label on top of it. **Fig. 2D** illustrates this behavior by representing the most active clone per pixel after running inference with a trained model on four MNIST test digits. This cloning mechanism allows us to represent the long-range interactions required for foreground-background segmentation using only pairwise factors. Our CMRF model is eight-connected and fully convolutional, meaning that there are only four different types of pairwise potentials connecting each pair of clones: vertical, horizontal, principal diagonal, and secondary diagonal (color-coded in **Fig. 2B**). It is important to note that these factors are learned during training from random initialization. This is akin to neurons developing contextual selectivity through visual experience, which we explain below.

In addition, pixel labels emit observed pixel intensities via emission factors (bottom vertical lines in **Fig. 2B**). In the noiseless case, these factors are deterministic: the `IN` and `OUT` pixel labels produce pixel intensity 0 (corresponding to the absence of a contour), whereas the remaining `CONTOUR` label produces pixel intensity 1 (corresponding to a contour). In practice, the observed binary images of pixel intensities are noisy (see **Fig. 2D**) with the emission factors being unknown. We propose to estimate them via maximum likelihood, which we discuss below.

Learning the full parameterization of this undirected eight-connected CMRF is a hard problem [40]. We therefore utilize query training (QT) [31], a recently devised method to turn a CMRF into a trainable inference neural network. During learning, we estimate the marginal probabilities of a random subset of pixel labels (the *query*) in the image given the other pixels labels, and minimize the cross-entropy of these marginals relative to the ground truth. The marginals of the query are estimated by unrolling parallel belief propagation (BP) [41], a local message-passing algorithm, over a small number of iterations. We refer the reader to [42, 31]for a deeper discussion of QT.

It is important to note the following features of our CMRF. First, in contrast with the simplified model in **Fig. 2B**, the models used in our experiments all consider one clone for the `IN` pixel label, one clone for the `OUT` pixel label, and 64 clones for the `CONTOUR` pixel label. We vary the number of clones of the `CONTOUR` in control experiments that demonstrate their role in higher-order representations. In addition, the CMRF takes as input a binary input of pixel intensities (left column, **Fig. 2C**), which is often set to be the output of an edge detector. Very roughly, this is akin to the input V2 gets from V1. During training, this noisy binary input is paired with a clean ground-truth foreground-background segmentation (right column, **Fig. 2C**).

We train a single CMRF for all of our experiments (with the exception being the ablation experiments in which we vary the number of clones and the top-down attention experiment, since in both cases we need to use a different model). This CMRF is trained on MNIST digits [43] that have been preprocessed to extract their contours and in which we have either a) added and removed edges^1^ (see **Fig. 2C** for a representative example); or b) removed some of the values, so that they effectively contain missing data (see **Fig. 4E**). Digit 8 is excluded from the training data, so that testing can proceed on an entirely new shape.

**Figure 3:**
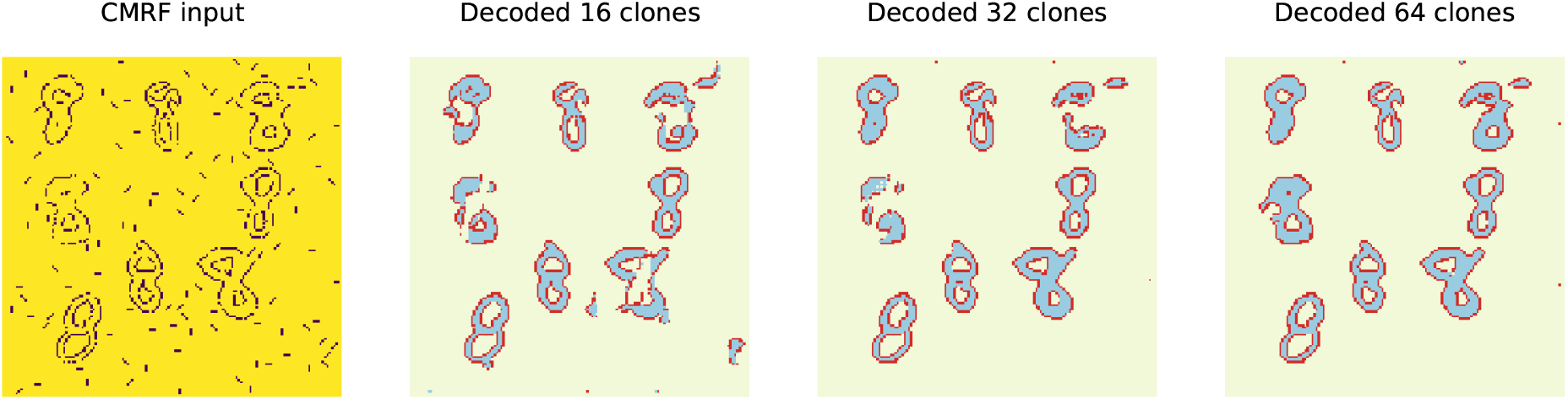
CMRF clones learn to encode higher-order contour dependencies. A CMRF with only 16 clones is unable to utilize the higher-order information to bridge large gaps. Models with more clones are able to better capture long-distance dependencies to improve their perceptual grouping performance

**Figure 4:**
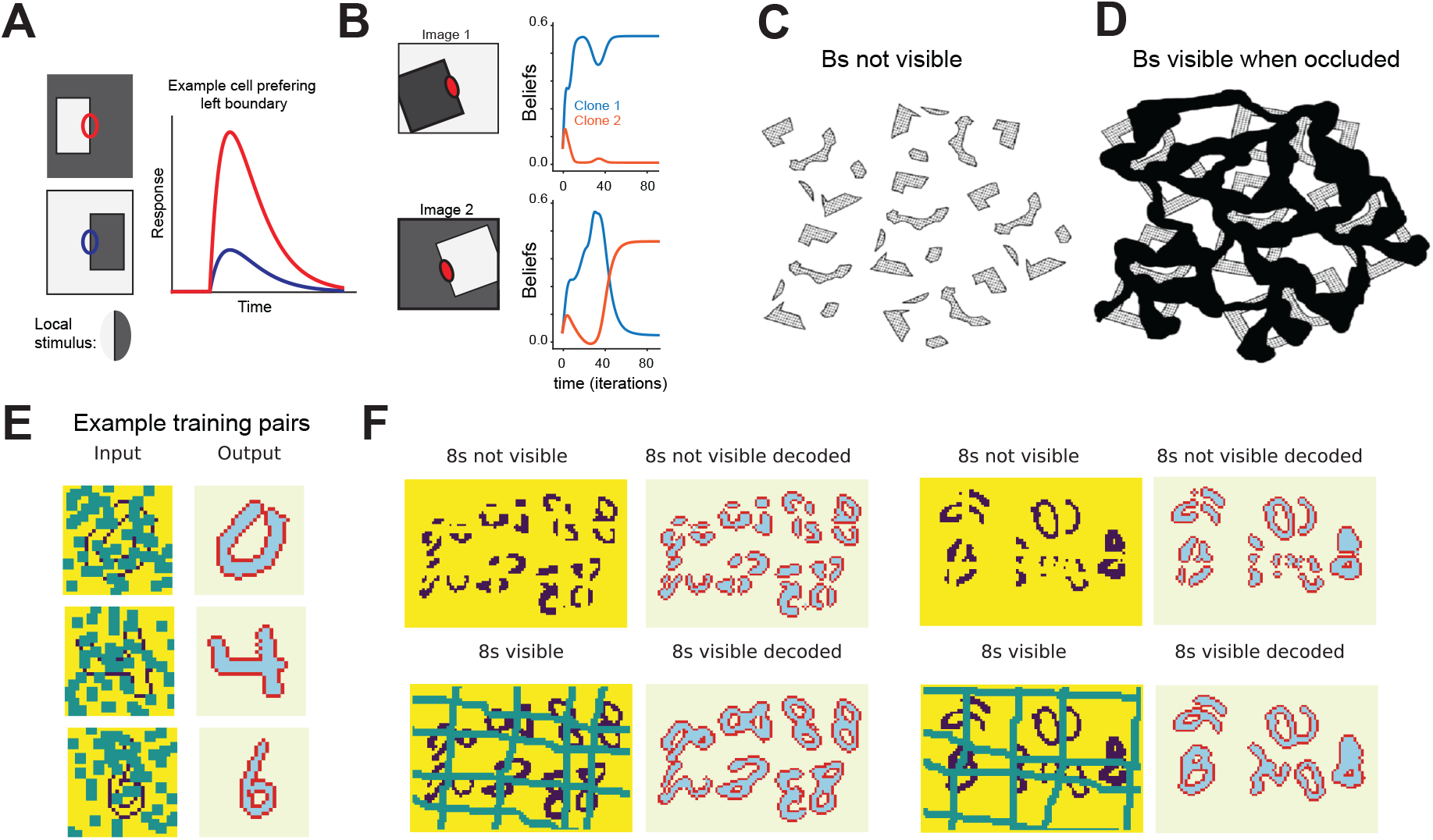
CMRF clones display border selectivity which allows it to complete surfaces and reason for occlusions. **A**: Schematic of the firing rate of a neuron that is selective for a border on the left. This neuron does not respond when the same border is on the right (blue) although within the receptive field (ellipse) the stimulus is the same. **B**: Temporal selectivity of two clones for the same edge. Similar to A, these clones also learn to prefer either a left-ward or a right-ward border. **C-D**: Bregman’s visual illusion. Although the fragments in C and D are the same, adding an occluder allows the characters to be recognized in D. **E** Input / ouput pairs of the second part of the training set. The input contains 3 × 3 patches of missing values, whereas the output is the ternary segmentation. QT can not only handle missing values as part of the training, but will also find parameters that are better suited for queries containing them. Clean binary inputs are associated with random block queries. The model is trained to recover the pixel labels segmentation. **F**: Experiment replicating Bregman’s Bs experiment using the MNIST digit 8. Without the occluding pixels, it is hard to perceptually group the digits. A CMRF trained on all MNIST digits but 8s is able to correctly identify the 8s when the occluder is present (top row) but not when it is absent (bottom row). See also Supplementary Materials **Fig. 1**.

During training, we use the foreground-background segmentation to derive the maximum likelihood estimates (MLE) of the pairwise potentials between pixel labels and pixel intensities, which are the bottom vertical lines in **Fig. 2B**. More precisely, these potentials correspond to the probabilities of emitting the pixel intensities given the pixel labels. As we know the exact segmentation at training time, we estimate these emission probabilities in closed form via maximum likelihood by computing the ratio of the empirical counts [41]. These estimates are then used for both training and testing. Finally, the potentials between pairs of clones are learned with QT [31] to recover the foreground-background segmentation from the noisy binary image. Subsequently, at test time, our CMRF model runs inference to estimate the marginal probabilities of all the clones in the image. It then computes the probabilities of each pixel label by summing over the probabilities of the clones corresponding to the same label. Each pixel in the test image is then assigned to the label with higher probability, that is, either an `IN`, `OUT` or `CONTOUR` label (**Fig. 2D**). We refer the reader to the [42, 31]for further detail on the CMRF training and inference algorithms.

How does the CMRF solve the border ownership problem? It has been proposed that assigning labels to the foreground and background requires integrating global context with local pixel information [8, 25]. Clones within our CMRF can specialize and learn higher order properties of the contour, such as its orientation and the side of the foreground on which they are located (cf. **Fig. 2D**). This is akin to border-ownership cells observed in the visual cortex that signal the foreground side of contours [8]. Therefore, in learning the contextual relationship between pixels, our CMRF is able to perform figure-ground segregation. We demonstrate this in the following sections through a series of carefully designed experiments.

Before going further, let us emphasize that all the results presented in **Fig. 2** and **Fig. 4** correspond to one CMRF model with *the same* set of parameters. As we mentioned above, the model has been trained with QT on a training set derived from the MNIST dataset, without 8s, composed of input / output pairs in the form of **Fig. 2C** and **Fig. 4E**.

### CMRFs can perceptually group and segment novel object categories

Our experiments show that the CMRF learns a generic prior of contours and surfaces that allows it to perceptually group novel object categories and segment them from the background despite noise, missing information, and clutter, and despite having only impoverished contour-only evidence. To demonstrate the ability of the CMRF to generalize to previously unseen object categories, we created novel categories of objects from MNIST digits by rotating them by 90, 180, or 270 degrees (cf. **Fig. 2D**). These rotated categories are never shown during training, and the model does not have any built-in rotational equivariances. CMRF achieves near-perfect recovery in noisy and cluttered settings, demonstratring that it learned a generic prior that allows this kind of recovery for entirely novel shapes. We quantify the model performance using the intersection-over-union (IoU, see Methods for more detail) score between the decoded digits and the ground truth, and report an average IoU of **92**.**19**% on this rotated MNIST test set of size 10, 000.

Since CMRF is fully convolutional, its perceptual grouping capability extends to scenes with multiple novel object categories and arbitrary image dimensions. In other words, despite being trained on a small 30 × 30 pixel image, it can segment a larger image of size 150 × 150. We demonstrate this in **Fig 2E**: we test our model on large images containing random upscaled 8s from the MNIST test set (all the 8s have been removed from the training set). These previously unseen digits are made 50% larger than the training images. We use the MLE emission probabilities from the original MNIST training set and reach an IoU performance of **91**.**65**% on 100 test images (see Supplementary Methods for more details). To illustrate the convergence of the model after a few iterations, we plot the temporal evolution of `IN` and `CONTOUR` beliefs for a random 0 from MNIST test set in **Fig. 2 G - H** and Supplementary Videos 1 and 2.

### Clones in CMRF learn to represent border-ownership and higher-order contours

Clones in the CMRF learn to represent border ownership, and exhibit inference dynamics similar to the border-ownership dynamics observed in neurobiology. A seminal neurophysiological study [8] showed that neurons exist in V1, V2 and V4, whose responses are side-of-figure selective even with identical local features in their classical receptive fields across displays. Consider the example shown in **Fig. 4A**. Locally, both the red and blue receptive fields see the same stimulus, which is an edge between white and gray. In response to the figure shown in the top panel of **Fig. 4A**, where the object (a white square) is present on the left side of the image, a (red) neuron preferring a left border would respond more than a (blue) neuron preferring a right border. Conversely, the blue neuron would respond more to the bottom image of **Fig. 4A**. Importantly, the low latency of these neural responses suggests that these border preferences can arise from a low-level mechanism without the delay of top-down signals from higher cortical areas. We replicated this experiment using the CMRF model. In **Fig. 4B**, we show the temporal evolution of beliefs in two clones corresponding to the edge indicated by the red ellipse. Similar to the neurons reported by **(author?)**[8], our clones are quick to respond to their preferences to figure either on the left (top, clone 1) or borders on the right (bottom, clone 2). Therefore, similar to neurons in V2, our clones develop side-of-figure preference that encodes ownership information at occlusion boundaries. It is important to note that the different copies of the same contour to represent different border onwerships emerge naturally in the CMRF when learning contour-surface-background factorization of images. This underscores the importance of these side-of-figure neurons in allowing the visual system to effectively reason border ownership.

In addition to border-ownership, different clones in the CMRF also serve to encode higher-order contours. We demonstrate this using three different CMRFs with 16, 32, and 64 clones each, but otherwise trained identically. When tested on the perceptual grouping task described earlier, the CMRF with fewer clones is unable follow object curvatures and bridge large gaps, leading to over-segmentation and spurious objects in the noise. As the number of clones are increased, the model is able to capture more higher-order correlations for better perceptual grouping **Fig. 3**.

### CMRF can use occlusion cues to bind disconnected object fragments into unitary percepts

An important demonstration of our ability to reason at occlusion boundaries is demonstrated in Bregman’s visual effect [16]. In this famous example, character B is not visible from isolated fragments shown in **Fig. 4C**. However, by simply adding a common occluder over the image, the very same fragments are easily recognizable as the complete character B shown in **Fig. 4D**. This demonstrates how unconscious perceptual mechanisms analyzing occlusion relationships reason for the occlusion (or try to figure out what is behind the occluder) and aid object recognition. This mechanism allows us to perceptually group the fragments into a more coherent percept.

To test the CMRF for the Bregman’s Bs effect, we created a novel “Bregman-8” test set that mimicked Bregman’s Bs using the MNIST digits 8s. **Fig. 4F** shows that the 8s are only recognizable for a human when occluding lines are present. This dataset falls naturally into our framework as occluding lines in **Fig. 4F** correspond to unknown test queries to the CMRF. Note that the digit 8 was never present in the training set of the CMRF. To solve this modified version of Bregman’s experiment, we do not perform any additional training and instead consider the same CMRF model used in the previous experiments, with the same parameters.

CMRF demonstrated the Bregman’s Bs perceptual effect when tested on the Bregman-8 test set. When tested on images without occlusion cue (top row, **Fig. 4G**), the CMRF fails to bind the different surfaces into unitary 8’s, and instead treats the disconnected fragments as different surfaces. When testeds on the corresponding images with occlusion cues (bottom row, **Fig. 4G**), inference in CMRF binds the disconnected segments into unitary percepts of 8’s, similar to the perceptual phenomena observed in people. Consequently, it is able to accurately identify at test time the 8s (unseen during training) when occlusions are present, reaching an IoU performance of **86**.**49**% on a test set of 100 images — as opposed to **74**.**92**% when no occlusions are present. See **Fig. 1** in the Supplementary Materials for more examples. CMRF offers an explanation for why the occlusion cue helps with binding disconnected objects – the occlusion cue acts as evidential explaining-away for the missing contours and surfaces, allowing probabilistic inference to give a higher score to grouping the disconnected segments.

### CMRF can use top-down attention to separate overlapping objects

Since border-ownership and object parsing are multi-stable, to parse challenging scenes with multiple novel overlapping objects, the CMRF representations need to be controllable via higher-order object priors. Several pieces of evidence suggest that top-down attention provides contextual information that is crucial for border ownership assignment [44]. In particular, one model posits that feedback from higher visual areas with larger, and hence more contextual receptive fields, endows V2 with contextual selectivity [45]. An example of top-down attention is provided in **Fig. 5A**. Depending on the focus of attention, one can either identify a vase or two faces in Rubin’s vase illusion.

**Figure 5:**
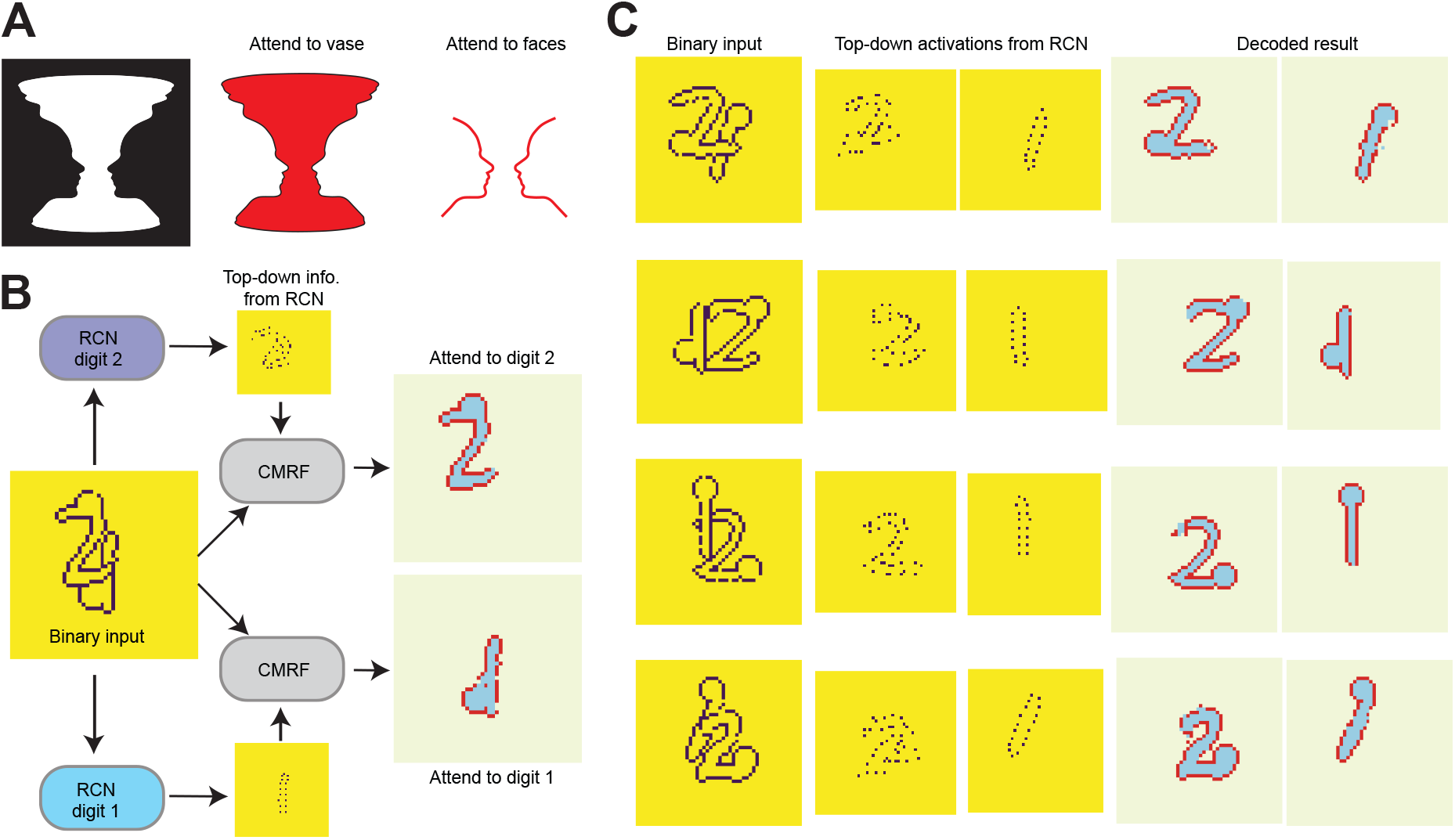
Top-down attention improves contour segmentation. **A**: Rubin’s vase experiment can be parsed into either a vase or two faces depending on the focus of attention. **B**: Implementation of attention modulation scheme. An RCN model is trained for each clean digit. The inference results of these RCN models are then used to inform the CMRF about which surface to segment. **C**: CMRF inference results on four different test images. For a given digit, the CMRF pairs the binary input with the top-down attention returned by RCN, and is trained to segment the digit from its background. See also Supplementary **Fig. 3** for additional results.

We seek to determine whether applying attentional modulation allows the CMRF model to segment different surfaces in an image. We created a new “overlapping MNIST” dataset by overlapping modified instances of digits 1 and 2. Each instance is created by drawing a random example from MNIST, computing its contour, attaching a circular perturbation to it, and deleting 5% of its pixels. The resulting “overlapping MNIST” dataset is shown in **Fig. 5C**. We split this dataset into a training and testing set of sizes 13, 500 and 1, 360, respectively. The task is to segment both digits, including the perturbation, from their background via inference. To address this task, we will combine our CMRF model (which has no notion of objects and only attempts to segment arbitrary surfaces) with a recursive cortical network (RCN^2^) model [37] (which is trained to detect individual digits with no perturbation). Note that none of these models can, by itself, solve the proposed task. The CMRF would try to discover a single surface (probably the digit in front) and delete the rest, whereas RCN would discover the presence of both digits, but because it only models clean digits, it would delete the circular perturbations, regarding them as noise. One could think of training RCN on perturbed digits, but the number of possible perturbations would produce a combinatorial explosion in the size of the training set. Instead, we let RCN model the global digit shapes and the CMRF the local, continuity-preserving distortions.

Our segmentation procedure is shown in **Fig. 5B**. It proceeds in two steps. In the first step, RCN is run to detect one of the two digits (“1” or “2”). This produces a binary *backtrace* of the “clean” digit (see **Fig. 5C** for examples), where any unmodeled perturbations (in particular, the circular ones) are removed. Then, this backtrace is provided to the CMRF as top-down information, softly encouraging the selected pixels to belong to the `CONTOUR` category. When the CMRF is run, it will combine the bottom-up evidence from the input image, the top-down evidence from the RCN backtrace, and the internal contour continuity bias to yield a clean ternary image in which a single digit is captured, the deleted pixels are recovered, and the circular perturbation is present as part of the digit’s surface.

In order for the CMRF to learn to integrate top-down and bottom-up evidence, we modify it to generate, in addition to the ternary image segmentation that was present in our previous experiments, the binary image that RCN outputs (the backtrace). This CMRF is then trained using QT on the “overlapping MNIST” dataset, augmented with the RCN backtraces^3^. Note that the same two overlapping digits appear twice in the QT training set; one associated with the RCN detection of the “1” and its ternary ground truth segmentation and another one similarly associated with the “2”. We use a single CMRF to obtain the segmentations of both the “1“ and the “2” in an image, but obviously, different RCN detectors, as depicted in **Fig. 5B**.

Examples of recovered digits from a separate test set are showin in the last two columns of **Fig. 2C**. Overall, with the top-down attention provided by RCN, our CMRF achieves an IoU score of **70**.**52**% for digit 1 and **82**.**43**% for digit 2. More test examples are displayed in Supplementary **Fig. 3**.

## Discussion

Parsing a visual scene with novel objects, binding disconnected parts of an object into a unitary percept, and determining which borders belong to which objects seem like a trivial tasks. However, it is unknown how the visual system is able to accomplish these feats, and no computational solution is currently known to approach the reliability of the primate visual system. In this paper, we proposed a neurobiologically inspired machine learning model, the CMRF, that tackles the computationally challenging problem of figure-ground segregation of unknown objects.

CMRF is built on the learnings from prior computational and conceptual models tackling this problem [25, 44]. Zhaoping Li’s model [25] demonstrated that lateral interactions, are sufficient to explain physiologically observed neural tuning to border ownership. However, the potentials involved in the model were not learned, and the model was tested only on stylistic scenes. In another model, von der Heydt and colleagues [44] rely on higher level areas that have larger receptive fields and modulate the activity in the lower level areas via back projections using grouping (G) cells. The G cells project back to the same cells they receive input from, facilitating their responses. Furthermore, O’Herron and von der Heydt [46] showed how this model could be extended to explain the remapping of border-ownership signals. Unlike Zhaoping’s model, the Craft model can explain the large context integration and the short latency of the border-ownership signals. However, as pointed out by [11], such strict template-based feedback does not allow the flexibility needed for tackling the shape variations of novel objects. The top-down attention control mechanism in the CMRF was inspired by the idea behind G cells, but goes beyond it because the top-down information is treated as a flexible prior that is then combined with bottom-up and lateral evidence. Moreover, CMRF also tackles the problem of occlusion inference in challenging settings. The difference in the surface apperance of objects and background provide a major cue for instance segmentation in typical computer vision benchmarks, but humans are able to segment objects even with just contour cues alone. Focusing on this surface-appearance-impoverished setting allowed us to focus on some of the core challenges in inferring border ownership [11].

CMRF provides an explanation for how border-ownership cells can emerge from learning a generic factorized contour-surface prior for objects, and its inference dynamics provides and explanation for the extra-classical receptive field properties of border-ownership cells. The emission structure of the CMRF was inspired by an observation from biology about clonally related neurons [30, 36]—-neurons that are genetically wired to share the same bottom-up receptive field. This structure turns out to be a crucial inductive bias that allows the CMRF to learn higher-order contextual representations using the clones. The clones learn to wire to different contexts [47], and some of these clones end up representing the global context of border-ownership. During inference, local evidence is propagated through a biologically plausible message-passing algorithm to come up with a globally coherent solution. Its emergent temporal dynamics of border-ownership signaling matches physiological observations. Altogether, we believe the CMRF is a first step towards a model of objectness compatible with neurobiological evidence.

CMRFs border-ownership and objectness representations are compatible with binding [3, 48]and top-down attention: in addition to being able to bind disconnected surface fragments into coherence, the resultant representation itself is amenable to top-down attention control [49, 50]so that further object properties like color, texture etc. can be dynamically bound [4] and indexed as needed. The top-down attention mechanism in the CMRF allows for higher-order object priors to be flexibly superposed such that novel overlapping objects can be segregated. Formation of object percepts is a fundamental step in cognition, concept-learning [4], and model-based object-oriented reinforcement learning [51], and the capability for dynamic binding could make the CMRF model suitable for tackling concept learning problems at the intersection of perception and cognition.

In addition, our work touches up on a central question in perception about how are feedforward, feedback, and lateral evidence coordinated in the visual cortex. Border-ownership is a challenging problem precisely because of the catch-22 nature of the problem where the ownership depends on the figure, while the figure itself depends on the borders and their ownership [11]. CMRF was designed by postulating computational roles for neurobiological observations about long-range cortico-cortical interactions in figure-ground segregation i[52, 53], separate processing dynamics for boundaries and surfaces [54], object-based top-down attention [19, 55], and clonal neurons [30]. By learning a model that is capable of solving several of the mid-level vision challenges [1] while being compatible with these observations, CMRF could provide an understanding of how the different neurobiological mechanisms work together.

Our work opens up several avenues for future research. In the current CMRF the surface model is simple, they just model and propagate whether a pixel is inside or outside an object. Perhaps, allowing more clones for the surface nodes, and forcing them to represent surface angles could learn a rich surface representation that includes surface curvatures, just like the higher-order contour clones. Furthermore, another set of surface nodes that represent surface appearance – shading, texture, etc. – could be learned jointly with the surface angle representation. Currently the top-down prior is based on known objects. By extending the CMRF hierarchically, a multi-scale representation of objectness priors could be learned, so that top-down attention can be based on entirely novel objects as well. In general, the CMRF can be thought of as a lateral layer in all levels of a multi-level generative model like RCN [50]. Problems like Bregman’s Bs and other perceptual completion problems have aspects of abstract reasoning – many of those are instances of unconscious inference. A hierarchical generative model that combines aspects of RCN and CMRF could act as the pereceptual symbol system [5] in a visual cognitive computer [4] for learning abstractions as cognitive programs.

## Methods

### Cloned Markov Random Field as a surface model

Our CMRF model is a bi-dimensional extension of the cloned hidden Markov model developed in the literature to model sequences of symbols [28]. Extending it to two dimensions naturally results in a cloned Markov random field (CMRF) presented in **Fig. 2B**. In our presentation in this work we have decoupled the hidden layer in two layers for ease of exposition (the pixel labels and the clones of each of them). But given that each pixel label variable (yellow circles in **Fig. 2B**) follows deterministically from the clones variables right on top of it (white circles in **Fig. 2B**), those two layers could be collapsed into a single layer with no change in the model. The (hidden) clone variables form an eight-connected grid and through the pixel label variables, each of them is connected to an observation, which is an image pixel. The clones can be grouped according to the pixel label, with each clone capturing different contexts of the observed variables. In **Fig. 2B**, the first 5 discrete values that a hidden variable can take deterministically result in the output variable (which is binary) being set to “contour”. That means that an observation of the type `CONTOUR` corresponds internally to 5 different “clones” in the hidden space, with potentially different internal meanings (such as different orientations or border ownership for that pixel).

### Query training for CMRF

The CMRF model is a eight-connected undirected probabilistic graphical model (PGM), which makes its learning notoriously hard due to (a) the difficulty of integrating out the hidden variables (the ”clones”) (b) the intractability of the partition function [40]. For directed models, a popular approach is to use variational autoencoders, but the encoder only amortizes inference for a single probabilistic query (new queries require separate training). In contrast here, we want our undirected CMRF to handle unseen queries without retraining.

We therefore introduce a systematic approach to turn a CMRF (and more generally any probabilistic graphical model) into a trainable inference neural network (NN). During training, for each image, we independently compute the marginals of a subset of *query* pixels’ clones given the other pixels’ clones, and minimize the cross-entropy between these estimated marginals and the ground truth. The *query* (subset of pixels) can be chosen uniformly at random for every image and every training iteration, or can be fixed when the evidence variables and the query variables are fixed. This approach is termed query training (QT) [42, 31].

Belief propagation (BP), a local message-passing algorithm for approximate marginal computation, appears as a natural approach to compute the marginals of the queried variables while using the same parameters for all queries. BP is scale-invariant: the computation of the partition function is not necessary since it does not affect its operation. BP is also known to converge to the exact solution in a finite number of iterations for tree PGMs [56]. We consider herein the loopy BP algorithm with parallel updates unrolled over a fixed set of iterations as our inference NN. We initialize our NN at random, using the the same set of parameters (weights) across all inference steps (layers). We minimize the cross-entropy with respect to the network parameters via stochastic gradient descent.

## Supplementary Materials

### Details of the experiment using occlusion cues to bind disconnected object fragments into a unitary percept

We describe here the experimental details corresponding to **Fig. 4** of the main paper.

#### Training set

We remove all 8 digits from the MNIST^4^ training set and keep the remaining 54, 000 entries. We pad each image and extract contours, obtaining 30×30 pixel images. Each entry in the dataset consists of a pairing of a contour images and the corresponding ground truth segmentation `CONTOUR-IN-OUT`. Two occluders (one vertical and one horizontal) are added to each training image (shown in green in **Fig. 4** and **Supplementary Fig. 1A**). Note that pixels covered by an occluder are treated as missing data (i.e., we do not know if the corresponding pixel is a `CONTOUR` or `NO CONTOUR`). The two occluders are: `RW-LR` from left to right and `RW-TD` from top to bottom. `RW-LR` starts randomly between the 8th and 18th pixel of the first columns and stops when it hits the last column. At each step, the probability of moving right is 0.9 and its probability of moving down is 0.1. We make the random walk 3 pixels high. Similarly, `RW-TD` starts randomly between the 12th and 22nd pixel of the first row, has a probability of moving down of 0.9 and left of 0.1 and stops when it hits the last row and is of width 3 pixels. Three examples are presented in **Supplementary Fig. 1A**.

#### Training procedure

We use a CMRF with 64 clones for `CONTOUR`, 1 for `IN` and 1 for `OUT`. We use QT and train for 150 epochs, unrolling the BP sum-product algorithm for *N* = 15 iterations. We use ADAM with a learning rate of 10^−2^ with minibatches of 24 images. In the training queries each pixel of the input image has a 0.5 probability of being part the corresponding random query. The emission probabilities of the CMRF are set to its maximum maximum likelihood estimates from training data, since we know both the pixel labels and the emitted image data. At test time we use the same setup, obtaining the bottom-up messages from each test image and running the sum-product algorithm for 15 BP iterations. This results in (conditional) marginal estimations at each variable, and in particular at the pixel labels. We assign each pixel to the label (`CONTOUR-IN-OUT`) with highest probability.

#### Metric

We evaluate our test performance with the intersection-over-union metric between the pixels with ground-truth labels `IN` or `CONTOUR` and the pixel labels estimated to be `IN` or `CONTOUR`.

#### Test set

Our test set consists of 20 test images of size 60 × 90. Each image has 4 vertical occluders and 4 horizontal occluders. Each occluder is generated by a random walk. The starting pixels are uniformly distributed and the transition probabilities are the same than for training. After creating the occluders, we loop through the MNIST test images of 8s—on which we randomly apply a 0, 90, 180 or 270 degrees rotation. Each digit is inserted in the free part of the image with largest intersection with the occluders.

Additionally, for each test image, we consider a variant in which the occluded pixels are set to be NO CONTOUR (instead of treated as unobserved, missing data). This is analogous to the Breigman’s Bs effect when no occlusion surface is present. We run the edge detector with the four filters shown in **Supplementary Fig. 4B** to obtain the CMRF binary inputs displayed in **Fig. 4E-F** also in **Supplementary Fig. 1B-C**.

**Supplementary Fig. 1:**
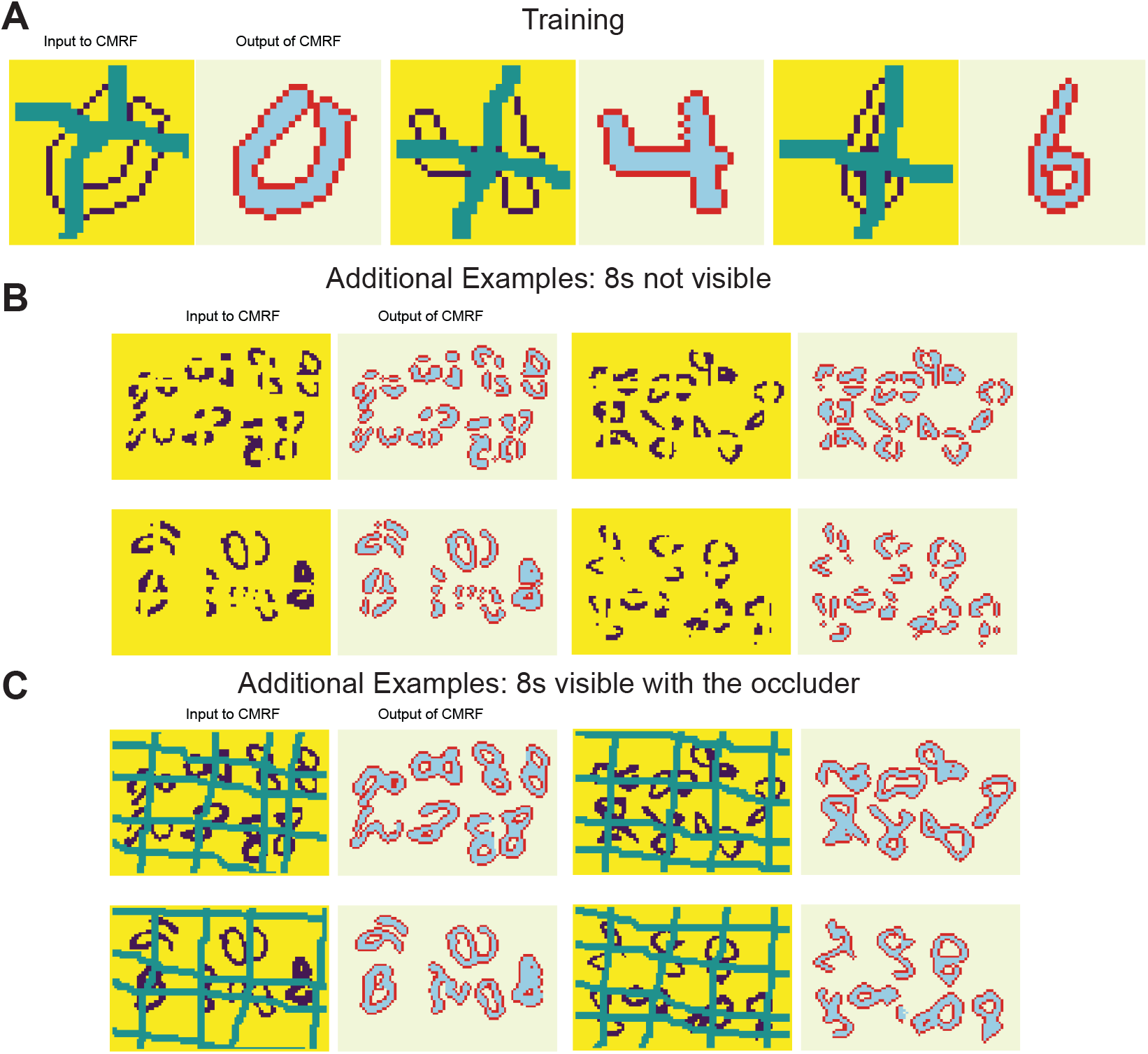
**A**: Examples of input-output training pairs for the second experiment. The CMRF is trained to recover the output segmentation from the binary image. The green lines used as occluder correspond to queries. **B - C**: A subset of eight test inputs, without (B) and with (C) occlusion, and their respective labels decoded by the CMRF for the second experiment.

### Analysis of the sensitivity to the number of clones

All the above experiments consider a CMRF with 64 clones for the `CONTOUR` state. Here we add results when training as above but using 16 and 32 clones instead. We consider the two generalizations described in the main text, i.e., rotations and upsampling. Table 1 shows the quantitative comparison and **Supplementary Fig. 2** shows the qualitative behavior. We observe that there is a performance degradation as the number of clones is reduced. This can be easily explained, since the reduced capacity limits the ability of the clones to capture the local context. One can expect that smaller contexts are captured and therefore more local, poorer reconstructions are produced.

**Table 1:**
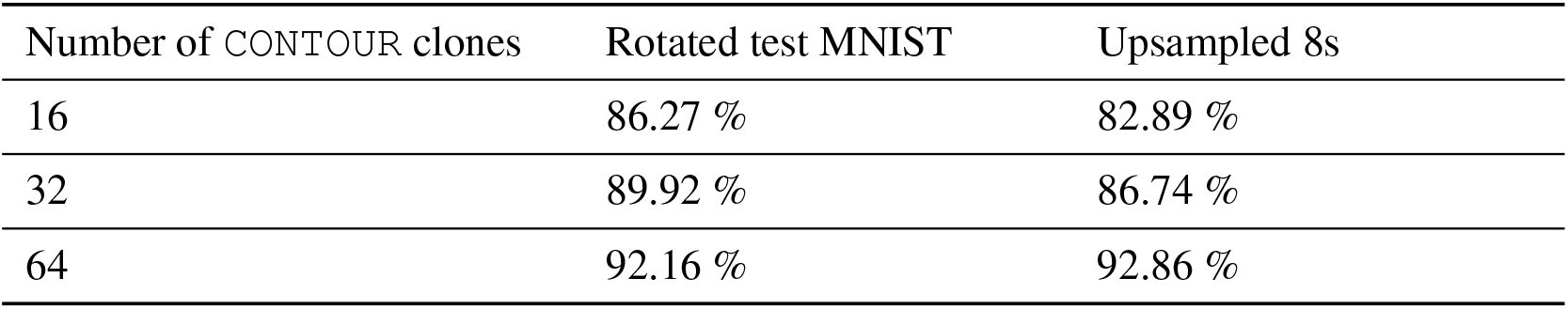
Performance comparison for three models trained with varying number of CONTOUR clones on the MNIST rotated test set and the upsampled 8s test set

| Number of CONTOUR clones | Rotated test MNIST | Upsampled 8s |
| --- | --- | --- |
| 16 | 86.27 % | 82.89 % |
| 32 | 89.92 % | 86.74 % |
| 64 | 92.16 % | 92.86 % |

**Supplementary Fig. 2:**
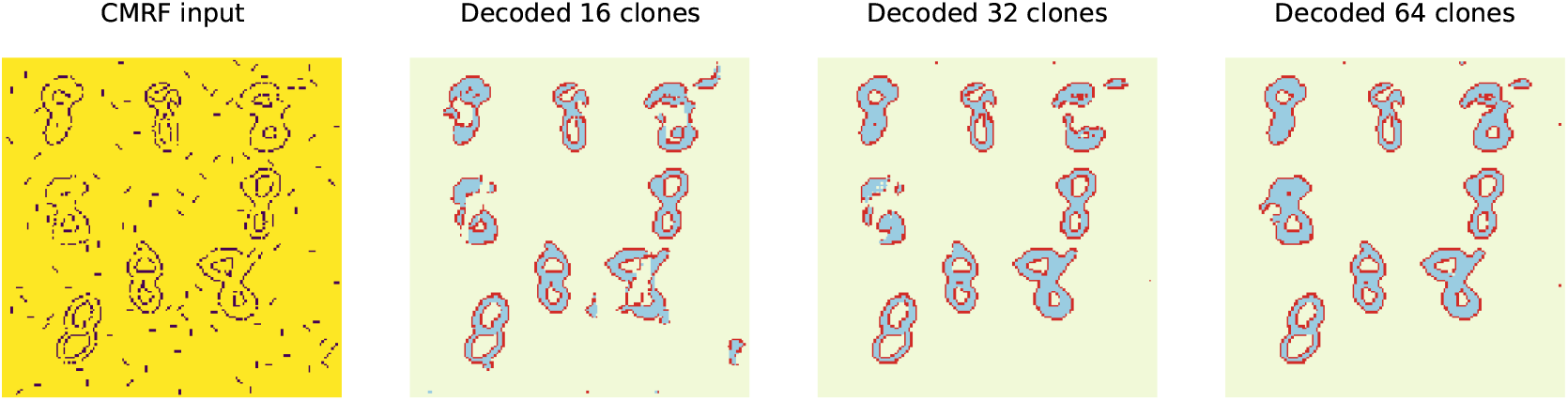
Decoding performance on the same image improves as the number of `CONTOUR` clones increase from 16 to 64.

### Details of the experiment using top-down attention to separate overlapping objects

We describe here the experimental details corresponding to **Fig. 5** of the main paper.

#### Training procedure

We first train two recursive cortical networks (RCN), introduced in [37], on the first 200 contour images with respective labels 1 and 2 of the MNIST training set.

We create a new “overlapping MNIST” dataset by overlapping pairs of training images containing a 1 and a 2. The resulting images have size 50 × 50. We consider the previously mentioned 200 pairs of 1s and 2s in RCN’s training sets. For each pair, we randomly select one digit to be on top and consider a fixed distance of 12 pixels and a uniformly selected random angle to shift one digit with respect to the other. We consider 100 variants of each training image, and for each of them we insert one circle of radius five on each digit at a random point along the contour, ensuring that the circles do not overlap and do not intersect with the other digit. Several images are presented in **Supplementary Fig. 3A**.

Finally, we run our edge detector to obtain contour images on which we run inference with the RCN models. For each digit, we consider the 30 highest-scoring detections returned by RCN inference, and keep the highest scoring detection such that at least 70% of its backtrace lies on the digit’s contour. We discard training instances where no such detection can be found for both digits. Notice that the discarding is done as part of the training procedure only. Each contour image produces two pairs of inputs to the CMRF, as displayed in **Supplementary Fig. 3B**. We use the first 10,000 images for CMRF training.

#### Test set

At test time, we only consider 1s and 2s from the MNIST test set. We create overlapping images similarly to **Supplementary Fig. 3A** and run the edge detector as above to derive the contour images. On each of the 1,360 test image, we run inference with both RCN models and keep the 30 highest-scoring detections returned by each model. Out of these 900 possible pairs, we retain the one for which the union of the backtraces maximizes the intersection-over-union with respect to the test contour image (not with the ground-truth, which is reserved for scoring), and use these backtraces as test input to the CMRF.

#### CMRF model

We consider a CMRF with 64 clones for `CONTOUR`, 1 for `IN` and 1 for `OUT`. We assume that, for each digit, the image with ground truth `CONTOUR-IN-OUT` segmentation independently emits the contour image (where both digits overlap) and the RCN backtrace of the corresponding digit. We set the emission probabilities in the model of the contour image and of the backtrace image given the pixel labels using the maximum likelihood estimators. We train for 10 epochs and use sum-product algorithm for *N* = 15 BP iterations, using ADAM with a learning rate of 10^−2^ with mini-batches of 50 images, and use the same bottom-up messages and number of BP iterations at test time.

### Demonstration of the CMRF model on non-digit objects

The contour-surface priors learned from MNIST digits generalize to objects that are not digits. We describe herein the experimental settings for our experiment on an object, presented in **Supplementary Fig. 4**. At test time, we consider an unseen *playa spray* object on a red background. We create 20 RGB test images, each of size 100 × 100. For each image, we iteratively place a uniformly selected in-plane rotation of the test object in a free part of the image. We stop when no such placement is possible.

We then run our edge detector as follows. We first convolve the four filters displayed in **Supplementary Fig. 4B** on the red channel of each test image and threshold the output: each filter returns a binary image which corresponds to contours activations above, to the left, down, or to the right, respectively. The edge detector returns the pixelwise maximum of these four binary activation images, and its output is then used as the CMRF input: contours are randomly missing and spurious edges randomly appear.

**Supplementary Fig. 3:**
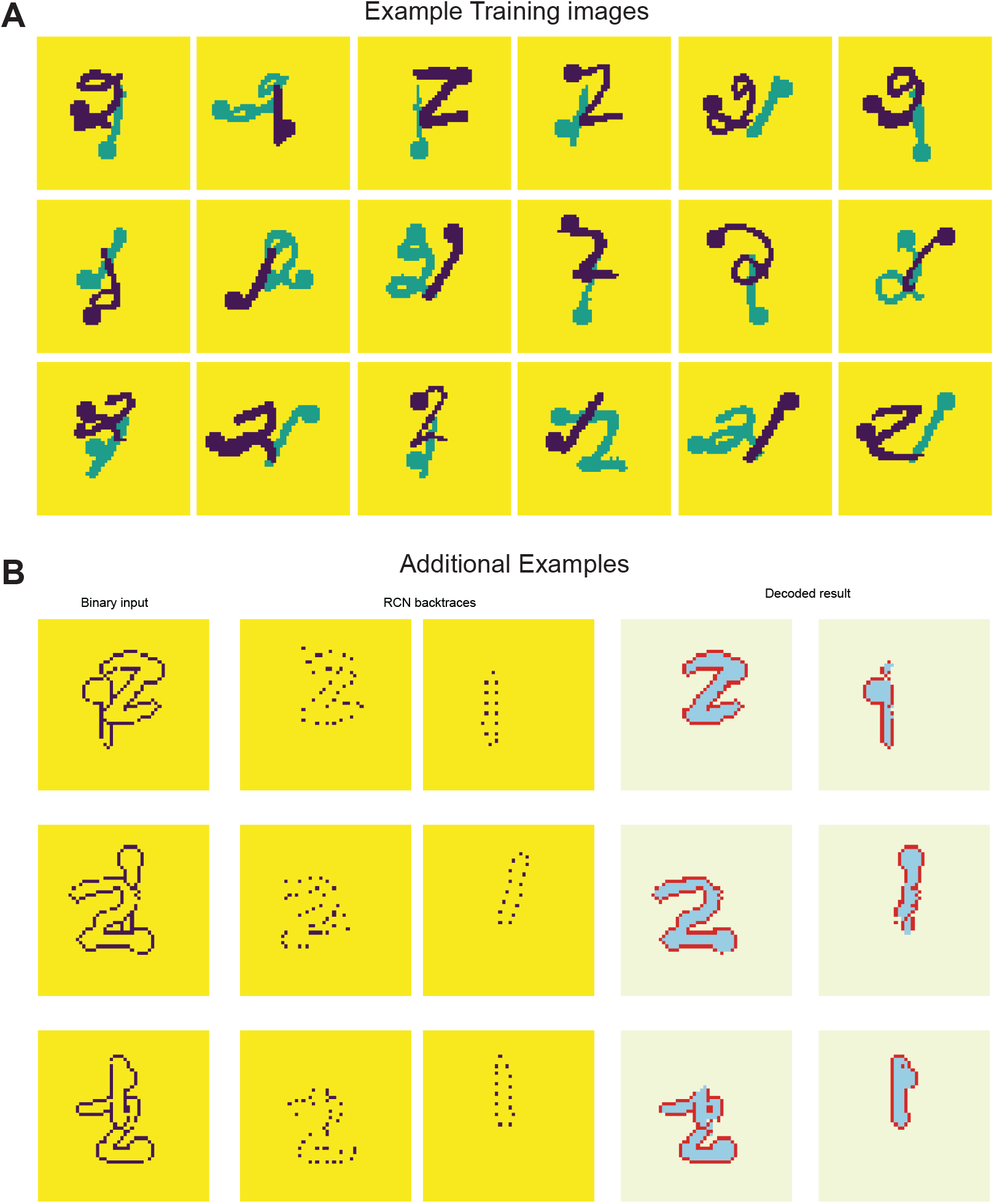
**A**: Examples of input-output training pairs for the third experiment. **B**: A subset of three contour test images and RCN backtraces for 1 and 2 digits, with their decoded CMRF labels, for the third experiment.

**Supplementary Fig. 4:**
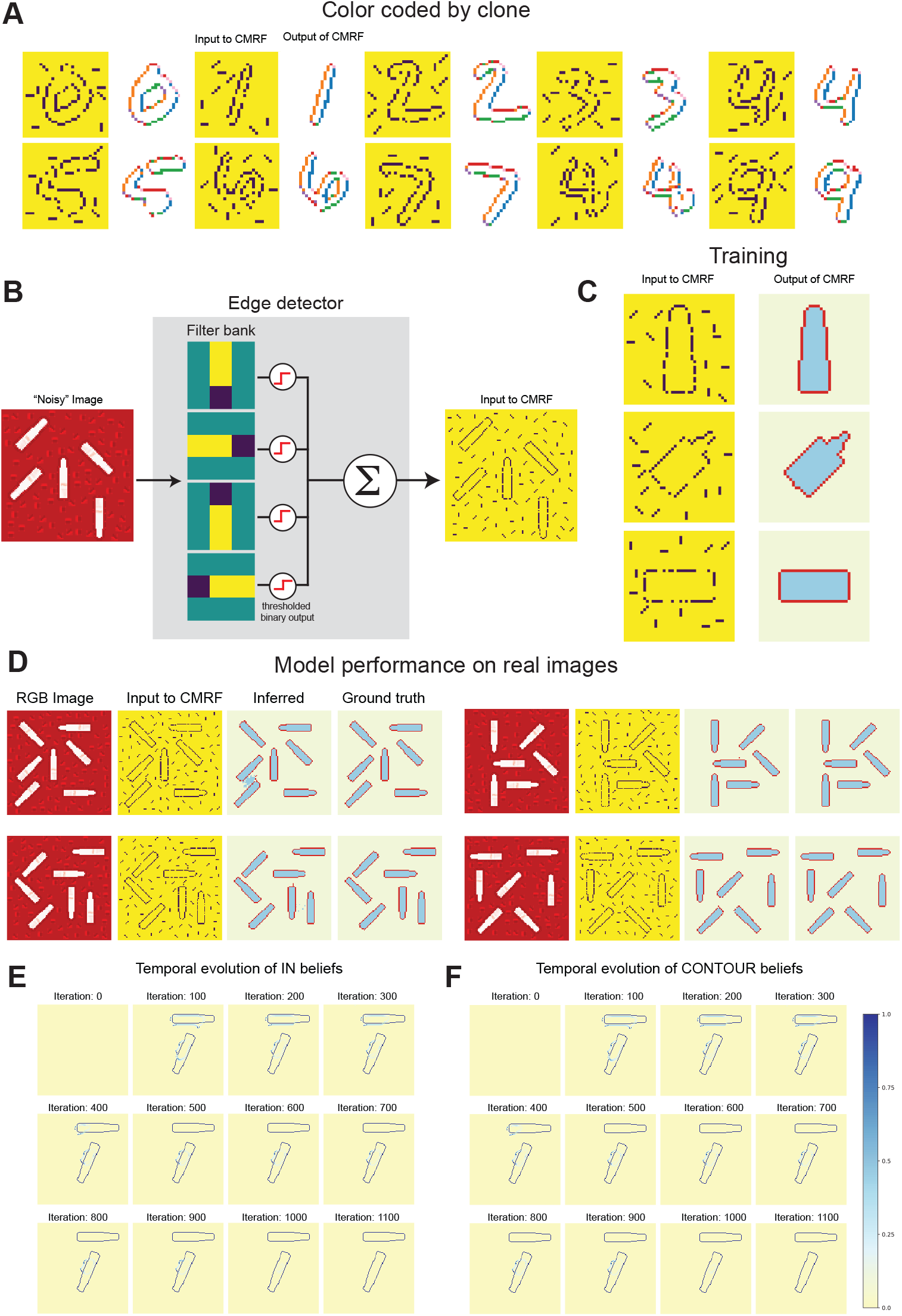
**A**: Each color corresponds to a different inferred hidden contour clone. The model has learned *with no supervision* to capture the local orientation of the pixel (which also reveals on which side the foreground is) as the best way to solve the denoising task. **B**: Schematic of edge detector. **C**: Examples of input-output pairs for the three objects used for the first experiment. The CMRF is trained to recover the output segmentation from the noisy binary images. Queries are selected at random: each pixel is selected with a 0.5 probability. **D**: A subset of five RGB input test pairs for the first experiment, with their respective CMRF inputs (edge detector output), labels decoded by the CMRF, and ground truth labels.

## Footnotes

1 More precisely, we add noise in the inputs of pixel intensities by (1) augmenting them with spurious edges at eight different orientations picked at random and (2) randomly switching off 20 percent of the pixels with intensities 1.

2 RCN is a structured generative PGM for visual parsing that systematically incorporates neurophysiological findings as inductive biases. This model has been shown to achieve state-of-the-art results on several vision benchmarks with greater data-efficiency compared to deep neural networks.

3 RCN is trained on clean, individual MNIST digits, with no circular perturbations.

4 The MNIST dataset can be found at http://yann.lecun.com/exdb/mnist/.

